# Chemometric Approaches for Developing Infrared Nanosensors to Image Anthracyclines

**DOI:** 10.1101/407650

**Authors:** Jackson Travis Del Bonis-O’Donnell, Rebecca Pinals, Sanghwa Jeong, Ami Thakrar, Russ Wolfinger, Markita Landry

**Affiliations:** Department of Chemical and Biomolecular Engineering, University of California, Berkeley, CA 94720; SAS Institute Inc. Cary, NC 27513; Department of Statistics, North Carolina State University, Raleigh, NC2 7695; Innovative Genomics Institute (IGI), Berkeley, CA 94720; California Institute for Quantitative Biosciences, QB3, University of California, Berkeley, CA 94720; Chan-Zuckerberg Biohub, San Francisco, CA 94158

## Abstract

Generation, identification, and validation of optical probes to image molecular targets in a biological milieu remains a challenge. Synthetic molecular recognition approaches leveraging the intrinsic near-infrared fluorescence of single-walled carbon nanotubes is a promising approach for chronic biochemical imaging in tissues. However, generation of nanosensors for selective imaging of molecular targets requires a heuristic approach. Here, we present a chemometric platform for rapidly screening libraries of candidate single-walled carbon nanotube nanosensors against biochemical analytes to quantify fluorescence response to small molecules including vitamins, neurotransmitters, and chemotherapeutics. We further show this approach can be leveraged to identify biochemical analytes that selectively modulate the intrinsic near-infrared fluorescence of candidate nanosensors. Chemometric analysis thus enables identification of nanosensor-analyte ‘hits’ and also nanosensor fluorescence signaling modalities such as wavelength-shifts that are optimal for translation to biological imaging. Through this approach, we identify and characterize a nanosensor for the chemotherapeutic anthracycline doxorubicin, which provides an up to 17 nm fluorescence red-shift and exhibits an 8 µM limit of detection, compatible with peak circulatory concentrations of doxorubicin common in therapeutic administration. We demonstrate selectivity of this nanosensor over dacarbazine, a chemotherapeutic commonly co-injected with DOX. Lastly, we demonstrate nanosensor tissue compatibility for imaging of doxorubicin in muscle tissue by incorporating nanosensors into the mouse hindlimb and measuring nanosensor response to exogenous DOX administration. Our results motivate chemometric approaches to nanosensor discovery for chronic imaging of drug partitioning into tissues and towards real-time monitoring of drug accumulation.

## Introduction

Availability of optical sensors for quantitative measurements of small molecules is a necessary component to effectively probe biological systems and advance our understanding of biological function, disease, and therapeutic efficacy. While the last several decades of sensor development have focused on organic and supramolecular chemistries for synthesizing new optical sensors,^1^ the recent emergence of single-walled carbon nanotubes (SWNTs) as nanoscale sensing elements has led to the development of numerous fluorescence-based chemical sensors selective for biologically-relevant molecular target analytes including reactive oxygen species, neurotransmitters, and proteins.^2–7^ The molecular specificity of SWNT-based nanosensors arises from unique configurations of a polymer-adsorbed phase, or corona phase, on the surface of the SWNT. Interactions between the polymer-phase and a target analyte modulate the innate fluorescence of the SWNT, producing an optical signal. Notably, the non-photobleaching fluorescence emission of these nanosensors falls in the short wavelength near-infrared (NIR) region (1000-1500 nm), where absorption and scattering by biological tissues is minimal, making them attractive candidates for long-term or chronic *in vivo* imaging of biological processes.^8,9^

The primary method of discovery for this class of SWNT-based probe has been through relatively low-throughput screening of candidate nanosensor libraries against potential target analytes of interest. However, SWNTs can be synthesized with up to 15 unique chiralities, each of which exhibits a unique fluorescence excitation and emission profile. Therefore, each SWNT chirality could form distinct polymer corona phases with unique molecular recognition elements, each with contrasting fluorescence responses to an analyte.^10^ The complex emission spectra of these samples and their chirality-dependent molecular recognition adds additional dimensionality to screening data. However, the most common approach to analyzing library screens has been monitoring changes to integrated fluorescence, losing valuable spectral information in the process. Although deconvolution of emission peaks and multivariate statistics have been used to monitor changes in peak intensity or solvatochromic shifts of known nanosensors,^11^ common chemometric approaches have yet to be implemented to analyze spectra obtained from initial library screens to identify new and unique analyte-nanosensor hits.

Here, we present chemometric approaches for analyzing fluorescence emission spectra obtained from screens of candidate SWNT nanosensors. Candidate nanosensors are each functionalized with a different polymer corona phase including polynucleotides, phospholipids, and amphiphilic heteropolymers, and screened against a panel of >40 biologically relevant analytes (see **Supporting Info** for analyte assignations). By implementing distance metric calculations, principal component analysis (PCA), and hierarchical clustering (HC), we demonstrate the rapid and quantitative identification of candidate analytes for nanosensors. These approaches uncover potential structure-response relationships between chemically similar analytes through identification and clustering of analytes that generate similar modulations to the SWNT fluorescence spectrum. We further demonstrate that this platform can be used to identify a novel NIR nanosensor for the detection of the chemotherapeutic doxorubicin (DOX) through a change in the nanosensor fluorescence intensity and up to a 17 nm red-shift upon exposure to DOX. Cardiomyopathy resulting from chemotherapies containing DOX is a primary reason for limiting its cumulative dose, impacting its efficacy as an anticancer treatment.^12,13^ Thus, monitoring DOX biodistribution in tissue using an optical nanosensor may help maximize the therapeutic value of DOX administration while minimizing its cardiotoxicity. As a first step to this end, we lastly demonstrate imaging of DOX in mouse hindlimb tissue with a DOX-responsive SWNT nanosensor, to confirm nanosensor functionality in a biological milieu. Our results motivate applications of this chemometric nanosensor discovery platform to generate SWNT nanosensors for biomolecule analytes, and their translation for use in biological tissues.

## Materials and Methods

### Candidate Nanosensor Synthesis

Soluble suspensions of SWNTs were prepared by combining between 0.2-5 mg of hydrated (5 wt% mixed-chirality SWNT (NanoIntegris) in deionized water) with 1 mL solutions of polymer or lipid. DNA samples contained 100 µM DNA oligo (Integrated DNA Technologies, Inc., standard desalting) in 100 mM NaCl. Phospholipid samples contained 2 mg/mL of 1,2-dipalmitoyl-sn-glycero-3-phosphoethanolamine-polyethylene glycol (2000) (DPPE-PEG, Avanti Polar Lipids, Inc.) in 1X phosphate buffered saline PBS and 0.2 mg SWNT. RITC-PEG-RITC polymer (rhodamine isothiocyanate-polyethylene(5000) glycol-rhodamine isothiocyanate) solutions contained 1 mg polymer, prepared using previously published methods,^3^ dissolved in 1X (PBS). Solutions were bath sonicated for 10 min followed by probe tip sonication using a 3 mm probe tip at 40-50% amplitude for 10 min. Samples were then centrifuged at 16100 cfg for 30 minutes to pellet insoluble bundles and contaminants and the supernatant containing product was collected. Concentration of SWNT was determined using sample absorbance at 632 nm and the extinction coefficient ε=0.036 µg/mL.

### Spectroscopy and Single Molecule Imaging

Emission spectra for SWNT nanosensors before and after exposure to small molecule analytes was collected using an SCT 320 spectrometer and PyLon-IR 1024/1.7 containing a liquid nitrogen cooled InGaAs array detector (Princeton Instruments). Nanosensors were excited using a beam expanded 721 nm diode pumped solid state laser (10-500 mW) coupled to an inverted microscope body (Axio Observer, Carl Zeiss) containing a custom filter cube set (800 nm SP FESH0800, 900 nm LP dichroic DMLP900R, 900 nm LP FELH900, ThorLabs) and motorized well plate stage (Marzhauser Wetzlar Gmbh & Co.). Emission light was collected and relayed to the slit of the spectrometer using plano convex lenses coated with antireflective coating AR-3 (ThorLabs). Single-nanosensor imaging experiments were captured using a NIRvana 640×512 InGaAs array camera (Princeton Instruments) and a 100x alpha plan-apochromat oil immersion objective with NA 1.46 (Zeiss). For surface-immobilized single molecule experiments, nanosensors were immobilized onto glass coverslip surfaces treated with (3-Aminopropyl)triethoxysilane (APTES, 10% v/v in ethanol, 2 min incubation followed by wash with PBS) by incubation for 3 min at 10 µg/mL concentration followed by a rinse with 1X PBS.

### Chemometric Analysis

Spectra obtained from fluorescence screens were normalized to the mean of the peak intensity for the (7,6) chirality SWNT of control measurements. Analysis was performed using JMP software suite (SAS Institute Inc.) and custom scripts written using MATLAB (R2017b, The Mathworks, Inc.) and its built-in functions for principal component analysis (`pca()`) and hierarchical clustering (`cluster(), linkage()`) with default parameters, unless otherwise noted. Curve fitting for DOX titration data was performed with the `fit()` function in MATLAB using a non-linear least squares method. The fluorescence intensity and peak wavelength of the nanosensor was assumed to be proportional to the fraction of binding sites occupied by DOX.^14^

### DOX Imaging in Mouse Tissue

Mouse hind leg muscle tissue was harvested following euthanasia and rapidly frozen in liquid nitrogen for preservation until use. SWNT nanosensor was loaded into tissue by incubation for 10 min in a 1X phosphate buffered saline (PBS) solution containing 10 mg/L nanosensor followed by thorough rinsing with 1X PBS. Images and emission spectra of the tissue submerged in a bath of 1X PBS were obtained using a 20x plan-apochromat objective with NA 0.8 (Zeiss). DOX was added to the bath to a final concentration of 100 µM and allowed to incubate for 10 min prior to collecting fluorescence spectra.

## Results and Discussion

### Chemometric Analysis of Simulated Nanosensor Fluorescence DATA

To evaluate different chemometric methods for analyzing SWNT candidate nanosensor fluorescence response data, we first constructed a simulated dataset representing expected results from a SWNT candidate nanosensor library screened against a chemical analyte library. The simulated dataset is comprised of 100 sample spectra each containing 4 Gaussian peaks to simulate the contributions of a sample containing a mixture of 4 different SWNT chiralities, comparable to the fluorescence emissions of (6,5), (10,2), (7,6), and (8,7) SWNTs. Should a candidate SWNT nanosensor respond to a particular analyte, we expect a change in the fluorescence emission spectra. These spectral changes can include (i) uniform fluorescence signal enhancement, (ii) uniform quenching, (iii) chirality-dependent enhancement and/or quenching, (iv) or red or blue shift in peak wavelength.^11^ As such, we included an example of each expected fluorescence response signal in our simulated dataset (**Figure 1A**): Uniform mild increase (*Response 1*), single-chirality increase (*Response 2*), uniform strong increase (*Response 3*), uniform wavelength shift (*Response 4*), no change (*Response 5*), asymmetric chirality-dependent change (*Response 6*), uniform decrease (*Response 7*), and single-chirality decrease (*Response 8*). Random noise was added to each simulated spectrum (0-5% total signal) along with systematic variance (0-10% total signal) corresponding to expected experiment-to-experiment variations. The completed simulated dataset is plotted as a heatmap, where color indicates intensity values (**Figure 1B**).

**Figure 1.**
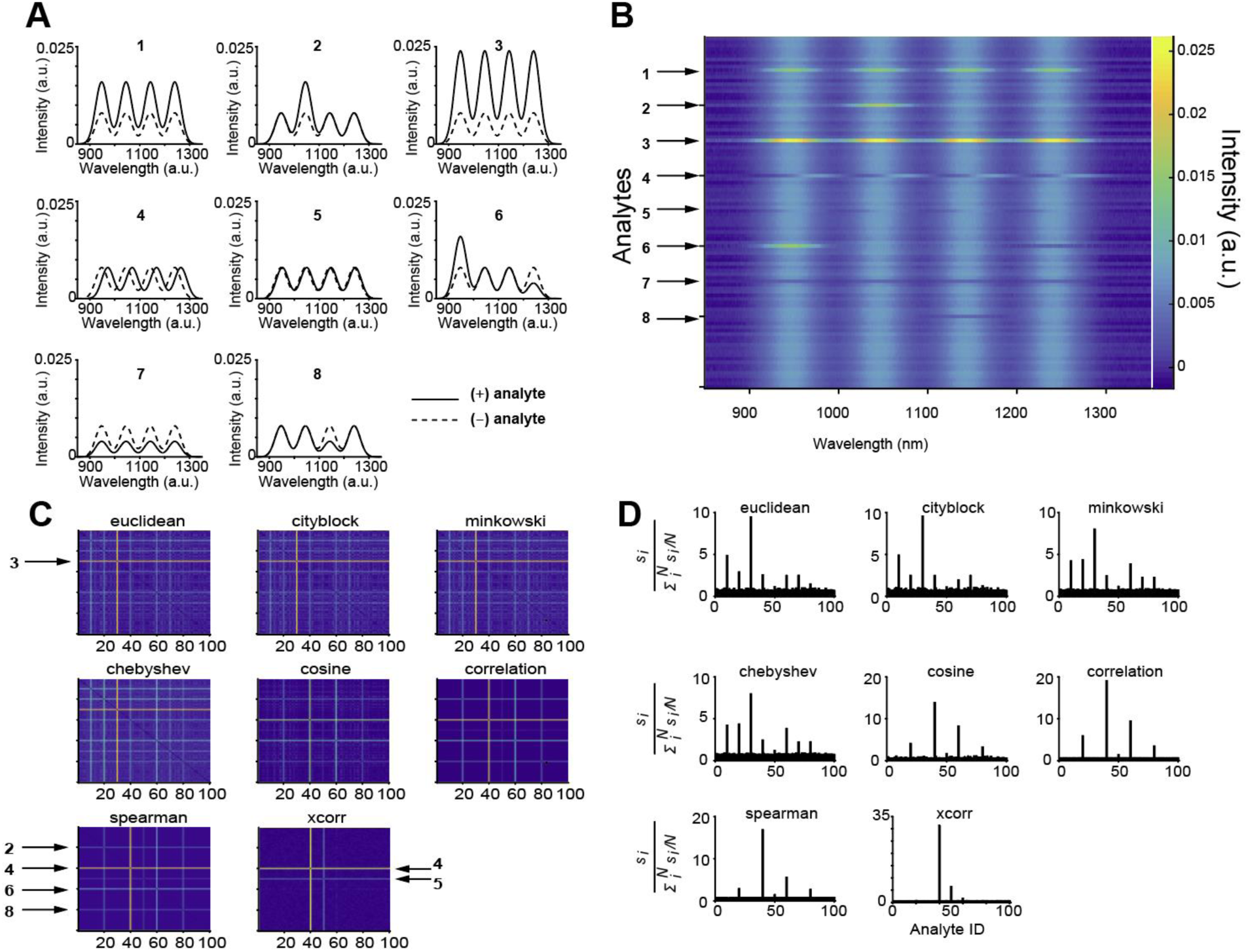
(A) Simulated emission spectra for SWNT nanosensor candidates after addition of different analytes. Dashed spectra correspond to original fluorescence emission. Solid lines correspond to fluorescence emission spectra of the nanosensor after analyte addition: Uniform mild increase (*Response 1*), single-chirality increase (*Response 2*), uniform strong increase (*Response 3*), uniform wavelength shift (*Response 4*), no change (*Response 5*), asymmetric chirality-dependent change (*Response 6*), uniform decrease (*Response 7*), and single-chirality decrease (*Response 8*). (B) Simulated library screen dataset containing 100 spectra displayed as a heatmap. Numbered arrows correspond to the analyte spectra shown in (A). (**C**) Pair-wise distance matrices for simulated library screens of SWNT nanosensor emission spectra. The choice of distance metric determines the analyte signals above noise and can be used to identify specific responses and candidate nanosensors. Arrowed numbers correspond to the simulated analyte responses shown in Figure 1a. (**D**) Response signals were calculated for each analyte using the different pairwise metric calculations. A potential signal hit or outlier is determined if the signal is greater than **3*σ*_*SD*_**.

With this simulated dataset, our goal is to explore statistical methods for identifying analytes that significantly and uniquely modulate the candidate SWNT nanosensor fluorescence emission, and to categorize analytes that induce similar responses. To start, we calculated commonly used pair-wise distance matrices using the following metrics: Euclidean, Cityblock, Chebyshev, Minkowski (order *p*=5), cosine, correlation, Spearman, and cross correlation (see **SI** for more information on distance calculations). Here, SWNT wavelengths and their corresponding intensities are treated as variables and values, respectively, while different analytes correspond to sample elements. **Figure 1C** plots pairwise distances calculated for our simulated dataset revealing that certain analytes were highlighted differently depending on the distance metric. We created a signal metric, s, by calculating the mean pairwise distance for each analyte, *s*_*i*_, and normalizing by the mean signal for all analytes, 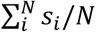. Of the nanosensor responses simulated, the response with a significant increase of fluorescence intensity (*Response 3*) yielded a large pairwise distance using the Minkowski metric family (Euclidean, Cityblock, and Chebyshev). Using the Chebyshev distance (Minkowski of order *p*=∞) as a signal metric, all responses, except for *Response 5*, produced a signal above a detection limit of 3*σ*_*SD*_, where the larger the fluorescence difference, the greater the distance metric value. Alternatively, using cosine, correlation, or Spearman distance for signal metrics, only *Response 2, 4, 6* and *8* were above the detection limit, with *Response 4* producing the largest signal. For these analyses, the magnitude of the fluorescence change dictated the strength of the calculated signal response. *Response 5*, which experiences a modest solvatochromic shift, failed to produce a signal above noise using any of the above distance metrics. Solvatochromic shifts are often preferred signals for use in complex biological media, since the wavelength shift readout can be more selective than a fluorescence intensity modulation.^3^ To overcome this limitation of identifying solvatochromic signals, we constructed a metric sensitive to wavelength shifts by calculating pair-wise cross correlations between spectra in a shifting reference frame (XCorr) to determine the wavelength offset. Using this metric, we obtained signals above noise for *Response 4* and *Response 5*, the only two simulated signals which corresponded to solvatochromic fluorescence shifts. Thus, results from our simulated nanosensor response libraries suggest that proper selection of a pairwise metric can be employed to readily screen for and quantify modulations in complex fluorescence emission spectra.

A powerful approach for rapidly analyzing spectra from library screens is principal component analysis (PCA), a multivariate technique employed to simplify complex chemical spectra and reveal underlying trends.^15–17^ Performing PCA on the simulated SWNT nanosensor spectra identifies an orthonormal set of variables (principal components) composed of linear combinations of the original variables (wavelengths) that maximize variance within the dataset. Subsequent components are chosen to maximize variance while remaining orthonormal to the previous components. A library of SWNT nanosensor response spectra can then be represented as vectors in a space comprised of the principal components that account for the most variance in the dataset. This simplification reduces dimensionality of the data while capturing the relevant changes to the height and position of spectral peaks without the computationally intensive cost of chirality deconvolution and peak fitting. **Figure 2A** outlines the results of performing PCA on the simulated SWNT nanosensor spectra dataset. Plotting each spectrum in principal component (PC) space demonstrates how this approach identifies outlying spectra. While most spectra are clustered around (0,0), spectra exhibiting a change in intensity or peak wavelength are distinctly separated. Each PC represents a particular characteristic of the spectra as represented by the component loadings. Analyzing the loading plots shows PC1 represents a uniform increase or decrease in intensity, PC2 represents a wavelength shift, and PC3 represents a non-uniform, or chirality-dependent, intensity change. Additionally, PCA identifies responses that are distinguishable from one another, a property leveraged previously for multiplexed sensing.^11^ For example, two nanosensors can be used simultaneously provided one undergoes a wavelength change, and the other an intensity change when bound to their respective analytes. Considering the simulated nanosensors in **Figure 2A**, a plot of the combined nanosensor spectra in (PC1, PC2) space can resolve and quantify the response of each nanosensor by projection onto its respective PC. This approach makes PCA particularly useful for screening nanosensor libraries to identify pairs for multiplexed sensing.

**Figure 2.**
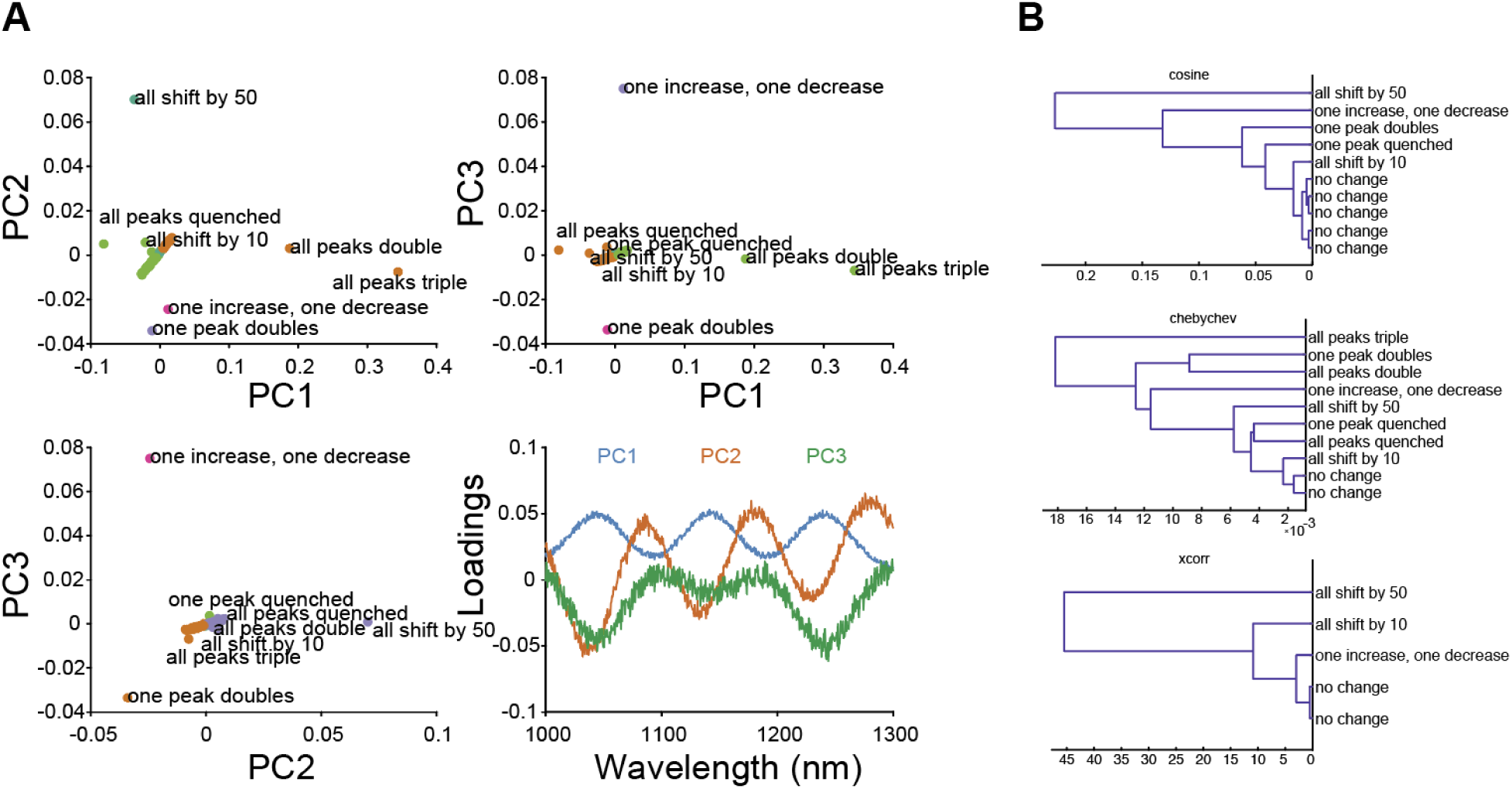
(**A**) Each simulated spectrum is plotted in principal component space of PC1, PC2, and PC3. A plot of the component loadings as a function of candidate SWNT nanosensor wavelength reveals the spectral charateristic for the first three principal componentents. (**B**) Hierarchical clustering dendrograms calculated with different pairwise distance metrics cluster similar analytes based on their modulation of SWNT fluorescence emission spectra.

Candidate nanosensor library screens can be further organized by clustering to identify analyte similarities. Hierarchical clustering (HC) assigns data into clusters based on the similarity of analyte modulation of SWNT fluorescence spectra. For example, in the simulated nanosensors library screen, HC is implemented to classify responses to different analytes into groups or clusters based on the similarity between spectra calculated using distance metrics. Groups are generated by merging similar spectra such that the increase in variance between members of the new group is minimized (Ward’s method). By performing these distance metric calculations, we clustered the simulated library screening data into groups of analytes that produced similar fluorescence responses (Error! Reference source not found.**C**). We chose to use the cosine, Chebyshev, and Xcorr metrics because each has different sensitivities to particular sensor responses: cosine is most sensitive for identifying signals where only a portion of the spectrum changes, Chebyshev for signals with large intensity changes, and Xcorr for spectra with peak wavelength shifts. Clustering analytes by the fluorescence response they induce on the SWNT nanosensor candidates provides insight into similarities between analyte properties and ultimately can inform mechanisms of interaction with the SWNT and its corona phase.

### Chemometric Analysis of DNA-, Polymer- and Phospholipid SWNT Nanosensor Libraries

To evaluate the efficacy of the above methods for curating information from real SWNT nanosensor candidate screening data, we prepared a library comprised of 10 SWNT nanosensors with distinct wrappings: 8 nanosensors synthesized with single stranded DNA oligo corona phases, 1 nanosensor with a heteropolymer corona, and 1 nanosensor with a phospholipid corona (see **Supporting Info** for details). Each SWNT nanosensor was screened against a panel of >40 biologically-relevant small molecule analytes including sugars, neurotransmitters, nucleosides, small molecule drugs, and vitamins (**Figure 3, Figure S1**). We included target analytes formerly shown to modulate SWNT fluorescence to ensure our approach could identify previously validated molecular analytes. These ‘known-responsive’ analytes were included as positive controls and include dopamine, ascorbic acid, thyroxine, melatonin, and riboflavin. Representative spectra compiled from DNA-corona nanosensor screens are shown in **Figure 3A**. The spectra are represented as heat maps, where each row represents a single SWNT fluorescence spectrum after exposure to an analyte, and color represents fluorescence intensity after normalization to the maximum intensity of the average control spectrum. Peaks in the emission spectra correspond to SWNT chiralities present in each sample. To identify analytes that modulated the SWNT nanosensor fluorescence, distance matrices were calculated using each distance metric (**Figure S1**) and subsequently used to calculate the fluorescence response signal, *s*, for each analyte (**Figure 3B**). Lastly, analyte “hits” (*s* > 3*σ*_*SD*_) calculated using each distance metric were compiled for SWNT nanosensors and visualized as a radial scatter plot, where spot size and radial position indicate the total number of “hits” for each analyte (**Figure 3C**). Results for all candidate nanosensors tested are presented in the **Supporting Information**.

**Figure 3.**
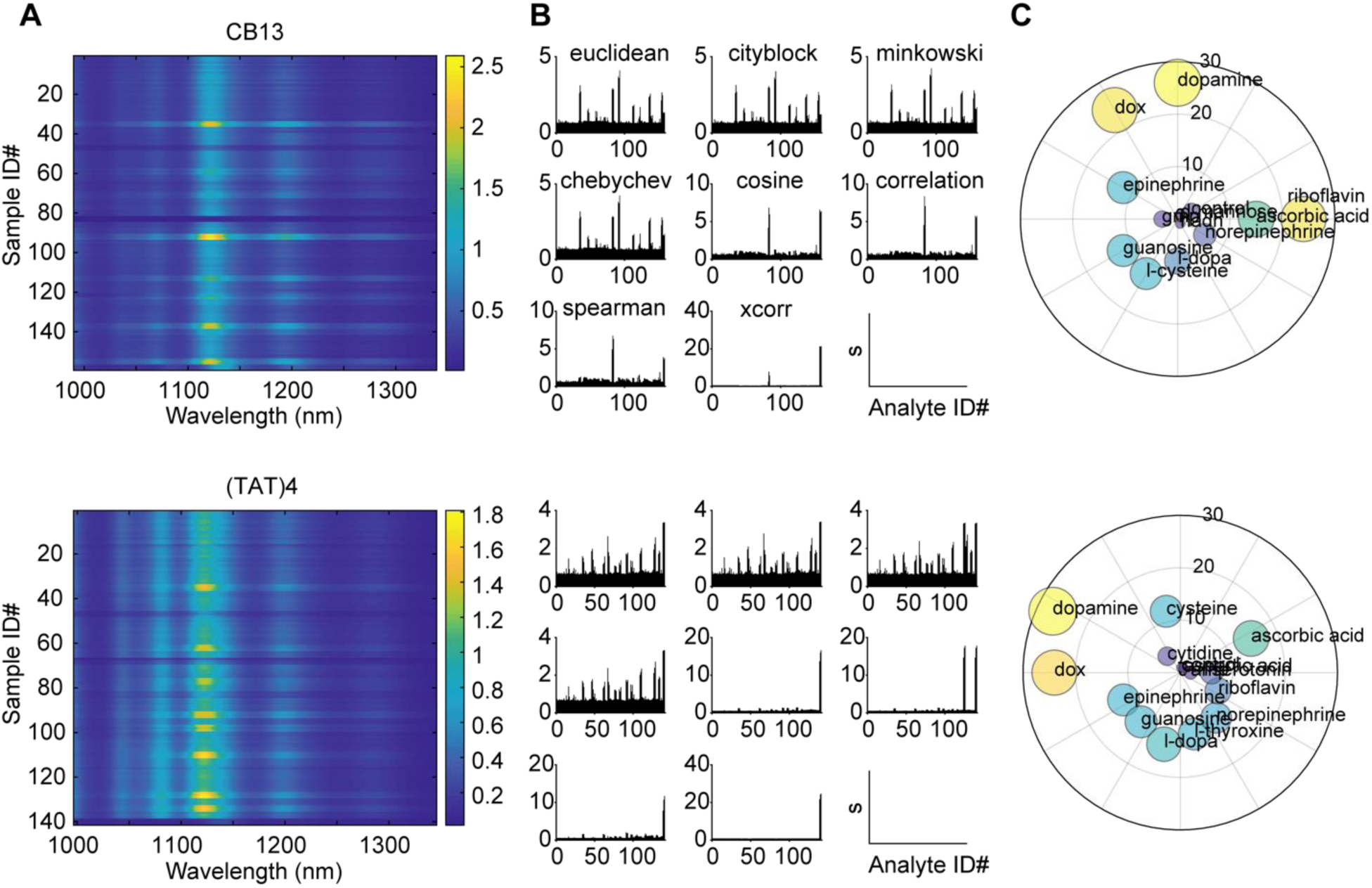
(**A**) Representative emission spectra heat maps for two DNA-SWNT nanosensor candidates upon exposure to an analyte library. Horizontal axes represent wavelength, vertical axes denote a particular analyte, and color represents normalized fluorescence intensity. Each analyte sample was screened in triplicate. (**B**) Response signal, *s* (vertical axes), for each spectrum (horizontal axes) calculated using different distance metrics. (**C**) Radial plots depicting analyte “hits” for the two SWNT nanosensors. The radial position and size of the circles scale with the total number of “hits” (*s >* **3***σ_**SD**_*) compiled from the combined results for all distance metrics.

For DNA-SWNT candidate nanosensor screens, we confirmed that previously known analytes elicit the predicted fluorescence responses on their respective SWNT nanosensors, such as the RITC-PEG-RITC heteropolymer-SWNT for which our screen confirmed a sensitivity to the hormone estradiol (**Figure S1**).^3^ Our approach also revealed new nanosensors with sensitivity to analytes guanosine, cysteine, and L-DOPA, which each induced a moderate fluorescence response to DNA-SWNT conjugates in a DNA sequence-specific manner (**Figure S1**). Additionally, we identify responses of several DNA-SWNT conjugates to analytes thyroxine and riboflavin, revealing that these analytes broadly impact the fluorescence of DNA-SWNT conjugates in a manner not specific to DNA sequence.^3^ We also identify novel responses of select DNA-SWNT conjugates toward other analytes in the library, including DOX and oxytocin. For instance, DOX was found to elicit strong fluorescence intensity and wavelength modulation when added to any of the DNA-SWNT conjugates. The response of phospholipid-PEG-wrapped SWNTs (**Figure S1**) was somewhat sensitive to DOX, riboflavin, and thyroxine. However, it was most sensitive to melatonin, consistent with previous accounts.^3^

The fluorescence response of candidate nanosensors to analytes from the experimental library was readily quantified with PCA (**Figure 4A, Figure S2**). The location of each point in PC space reflects both the magnitude and nature of the fluorescence response. Plotting the library screening data in principal component space depicts strong and/or unique analyte responses as outlying points. A majority of fluorescence responses are clustered around (0,0) in PC space, indicative of negligible candidate nanosensor fluorescence response, while responses that fall further from the origin strongly and/or uniquely modulated the SWNT fluorescence. One such ‘hit’ is shown in **Figure 4A** depicting representative data for DNA-SWNT nanosensor CB13. The nature of the response can be inferred from the plot of principal component loadings (**Figure 4A**): PC1 represents an increase in fluorescence emission intensity with longer wavelength SWNT chiralities showing a larger relative increase in emission than shorter wavelength emitting SWNT chiralities. Analytes with large PC1 scores include dopamine, ascorbic acid, and norepinephrine, consistent with previous observations.^14^ PC2 and PC3 represent fluorescence responses via both fluorescence intensity and wavelength shifts. Both DOX (Cluster 6) and riboflavin (Cluster 5) fall furthest from the plot origins for both PC2 and PC3 axes, consistent with their observed responses of quenching and shifting fluorescence emission.

**Figure 4.**
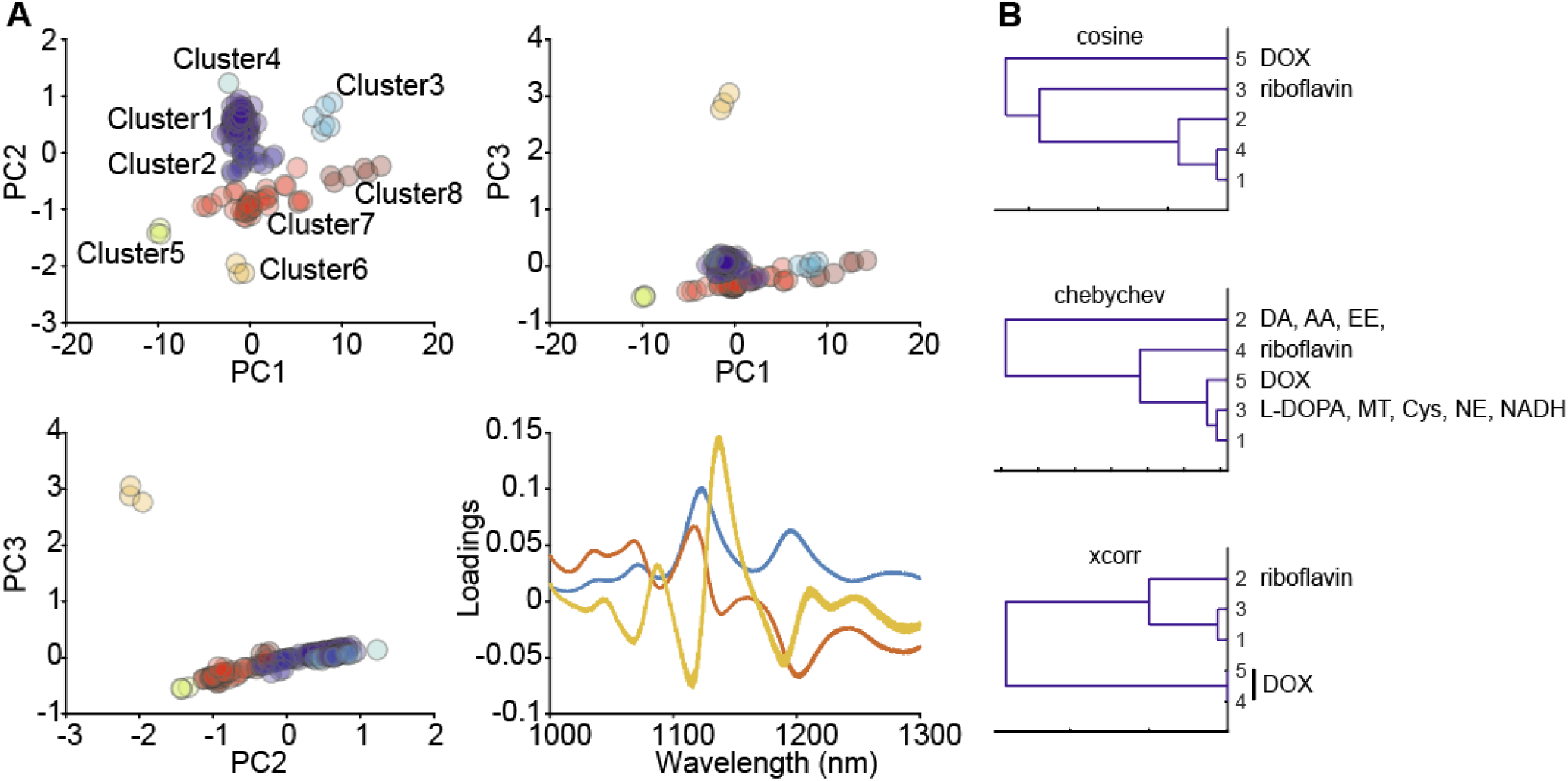
(A) Principal component analysis for an analyte library screened against the DNA-wrapped SWNT nanosensor CB13. Each dataset can be plotted using different principal component coordinates. Samples are clustered using hierarchical clustering computed with centroid distances calculated using a Euclidean metric and normalized principal components. Cluster assignments for each sample are listed in the Supporting Information. A plot of the loadings represents the spectral characteristic of each principal component. (B) Hierarchical clustering dendrograms generated using three different distance metrics for library data acquired for the DNA-wrapped SWNT nanosensor CB13. DA: dopamine, AA: ascorbic acid, EE: epinephrine, DOX: DOX, NE: norepinephrine, Cy: L-cysteine.

To investigate the relationship between fluorescence modulation of different analytes eliciting similar fluorescence responses, we next performed hierarchical clustering on the spectra library using cosine, Chebyshev, and Xcorr metrics. Representative data for the DNA-wrapped SWNT nanosensor CB13 is shown in **Figure 4B**. Applying the cosine and Xcorr metrics, we observe that all samples containing DOX and riboflavin were clustered separately from the rest of the analyte library. Both DOX and riboflavin induced a change in fluorescence emission intensity as well as a peak wavelength shift. When applying the Chebyshev metric, both DOX and riboflavin separate into a third distinct cluster. Additionally, samples containing dopamine, epinephrine, and ascorbic acid are contained in a single cluster, while L-DOPA, melatonin, L-cysteine, norepinephrine, and NADH are clustered separately. We next rationalized the observed clustering with prior reports of SWNT fluorescence modulation by different classes of analytes. The ability of catecholamines (dopamine and epinephrine) and ascorbic acid to brighten the fluorescence of DNA-SWNTs is attributed to their redox activity, however, their responses cannot be explained by redox potentials alone.^18^ L-DOPA, melatonin, and norepinephrine, all of which have an aromatic diol and similar redox chemistry,^19^ are more polarizable and contain an additional rotatable bond compared to dopamine, epinephrine, and ascorbic acid. This suggests a combination of polarizability and molecular bond flexibility properties may account for the differences in the fluorescence response of DNA-SWNT. Conversely, DOX and riboflavin, in addition to being oxidizing and light absorbing, both contain large planar aromatic groups. These chemical properties may contribute to increased steric and pi electronic interactions that distinguish the optical response induced on SWNT by DOX and riboflavin over the other analytes investigated. These analyte clustering trends were similar for the other DNA-wrapped SWNTs examined, with some clusters also containing analytes found to moderately modulate SWNT fluorescence, including glutamic acid, estradiol, oxytocin, serotonin, ampicillin, creatine, and DOPAC (**Figure S3**). As expected, analytes screened using the polymer-wrapped and phospholipid wrapped SWNTs likewise clustered according to their impact on SWNT fluorescence modulation. Furthermore, HC was also applied to analytes in PC space (**Figure 4A**) to identify distinct clusters far from (0,0), aiding in the identification of nanosensors for multiplexing.

### Evaluating SWNT Nanosensors DOX Detection

The above chemometric analyses revealed several unique DNA-SWNTs for implementation as nanosensors in the selective optical imaging of DOX. Direct in-tissue imaging of DOX during chemotherapy can help personalize and monitor dosage levels to minimize damage to cardiac tissue while maximizing therapeutic efficacy. Towards this end, we compared the DNA-SWNTs identified in analyses above as responsive to DOX and evaluated their responses for use in biological tissue. We first compared 12 distinct SWNT nanosensors wrapped in different DNA sequences to understand the sequence-dependence of DOX recognition and SWNT nanosensor fluorescence modulation. **Figure S1** shows that nearly all DNA-SWNTs exposed to 100 µM DOX experienced a quenching of fluorescence emission intensity. Additionally, DNA-SWNTs exhibiting the greatest degree of quenching also experienced up to an 8 nm red-shift in peak emission intensity. Curiously, the fluorescence emission of CB4 and CB14 SWNTs were not modulated by DOX despite having both a similar nucleobase sequence and spectral shape to responsive constructs, such as CB13. A more detailed comparison of each DNA-SWNT preparation revealed a weak correlation between DOX response, AT composition, and DNA sequence affinity for the SWNT, as measured by suspension yield (**Figure S4**). These results suggest SWNT surface coverage density by DNA plays a role in the conjugate’s interaction with DOX, and that differences in nucleobase adsorption energy to SWNTs renders certain ssDNA sequences more or less prone to fluorescence modulation by DOX.^20,21^ Furthermore, this analysis revealed that CB13-SWNTs, which experience both a quench and a shift in fluorescence upon exposure to DOX, was the nanosensor construct with the most robust change to its fluorescence emission, making it an optimal candidate for DOX detection.

We next investigated the role of DNA coverage density and nanosensor response to DOX by synthesizing CB13-DNA-SWNT constructs with variable DNA surface densities. By changing the SWNT:DNA mass ratio from 1:0.8 to 3:0.8 to 5:0.8, we observe a change in both the emission spectra of the CB13-DNA-SWNT constructs, as well as differences in the fluorescence response to DOX (**Figure 5A**). As the SWNT:DNA mass ratio was increased, the baseline intensity of the longer wavelength (larger diameter) SWNT chiralities decreased. Additionally, larger-diameter SWNTs experienced a sharp increase in fluorescence emission when exposed to DOX, in contrast to the decrease in emission intensity observed for nanosensors prepared using a lower SWNT:DNA ratio corona. We posit that decreasing the DNA to SWNT mass ratio causes an increased DNA wrapping pitch and exposes a larger proportion of the SWNT surface to the aqueous phase and thus susceptible to fluorescence quenching by oxygen, influencing SWNT baseline fluorescence emission intensity in a chirality-dependent manner.^22–24^ In this model, DOX binding to the SWNT via hydrophobic and pi-pi stacking interactions ^25^ would displace water, protons, and oxygen from the SWNT surface, reversing defect mediated quenching.^24^ Curiously, displacement of solvent from the SWNT surface, and the subsequent change in dielectric constant, is expected to induce a blue-shift in fluorescence emission, rather than the red-shift we observe.^26^ We therefore propose an alternative mechanism, whereby DOX modulates nucleobase stacking on the nanotube surface, inducing a conformational change in the DNA. Increased accessibility of solvent to the SWNT surface induces a red shift in fluorescence emission,^27,28^ while changes to the periodic surface potential on the nanotube surface modulate exciton lifetimes, which results in an increase in fluorescence emission.^29^ This increase in fluorescence is likely dependent on the formation of a highly ordered arrangement of the polymer on the SWNT surface, corroborated by the dependence on DNA surface coverage density we observe. These hypotheses are further corroborated by single-molecule microscopy measurements of individual nanosensors (**Figure S6**) to confirm that the fluorescence intensity changes are reversible, suggesting a transient, non-covalent interaction with DOX.

**Figure 5.**
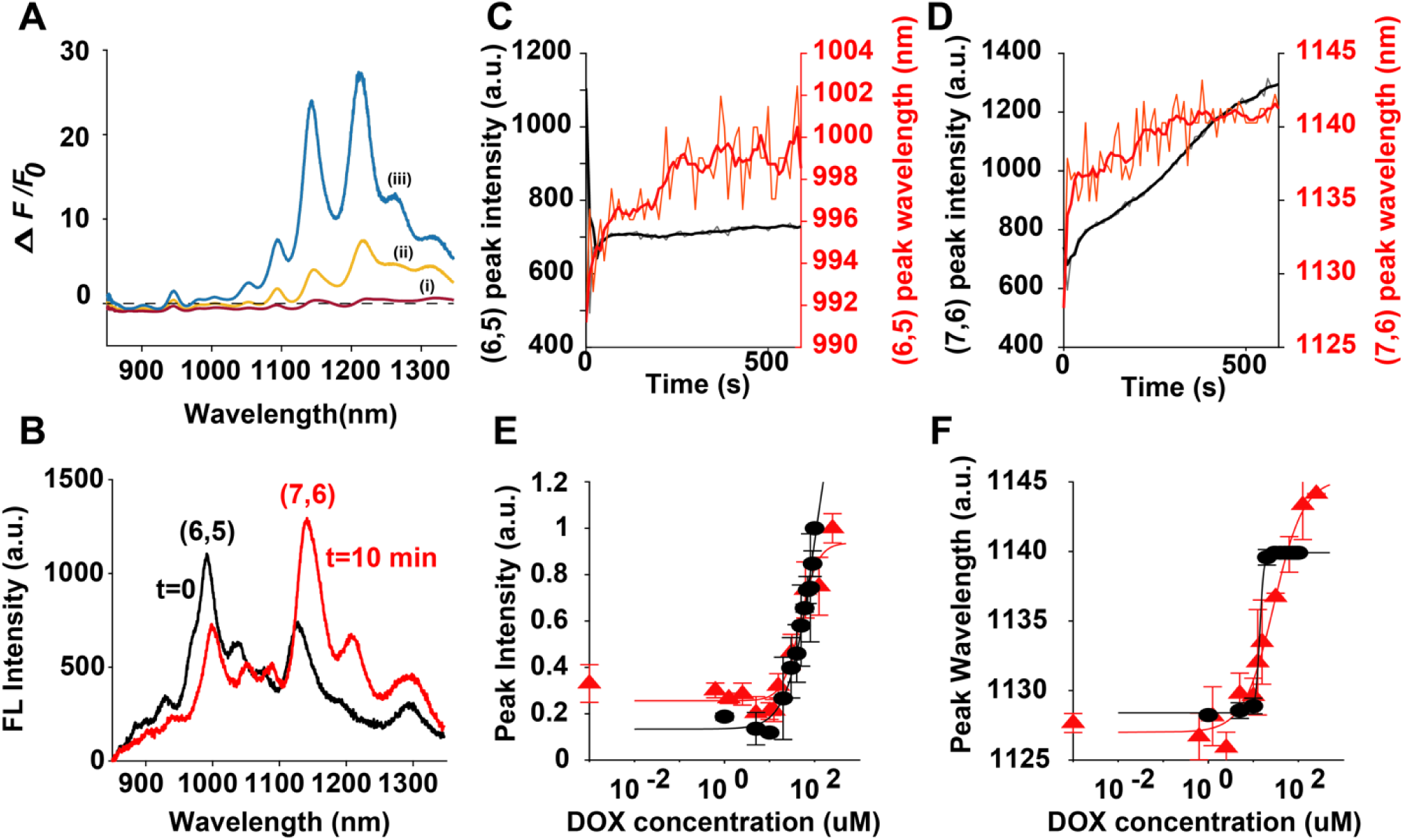
(**A**) Fluorescence response (*F-F*_0_/*F*_0_) for spectra of CB13 DNA-SWNTs upon addition of 100 µM DOX. Changing the ratio of SWNT:DNA (w/w) alters both the fluorescence emission intensity and response to DOX. 1 mL samples of 100 µM DNA suspending (i) 1 mg (red), (ii) 3 mg (yellow), or (iii) 5 mg (blue) SWNT. Each spectrum represents the average of 3 technical replicate measurements. (**B**) Fluorescence emission spectra of CB13-DNA-SWNT before addition of (t=0), and after 10 min incubation with (t=10 min) 100 µM DOX. Time series traces of the (**C**) (6,5) and (**D**) (7,6) peaks reveal that the peak wavelength (red) shifts continuously over 10 min while the peak intensity (black) for (6,5) plateaus rapidly compared to (7,6). (**E, F**) Fluorescence response of normalized intensity (**E**) and wavelength (**F**) for the (7,6) chirality peak of CB-13-DNA-SWNT versus DOX concentration. Triangles (red) and circles (black) represent separate experimental replicates. Each point is the average of 3 technical replicates with standard deviation as error bars. Curve fits were calculated using non-linear least squares.

Owing to the robustness of a turn-on fluorescent nanosensor compared to a turn-off readout, we further characterized the CB13-DNA-SWNT nanosensor using the 3mg mass ratio, which exhibited the largest intensity increase in response to DOX. Time-resolved measurements of the fluorescence response to DOX reveals a chirality-dependent fluorescence response of the mixed chirality CB13-DNA-SWNT sample (**Figure 5C-D**). (7,6)-chirality SWNTs experience an intensity increase and 12 nm red-shift that reaches steady-state after 10 min incubation with DOX. In contrast, DOX induces an 8 nm red-shift and quenches the fluorescence of (6,5)-chirality SWNTs by 30% and stabilizes within seconds. Using the (7,6)-SWNT emission peak intensity and wavelength, we constructed DOX concentration calibration curves for the CB13-SWNT nanosensor response (**Figure 5E,F**). Intensity and peak wavelength curves are fit to the Hill equation, used previously to characterize the kinetics of SWNT nanosensors with their analytes.^14^ From Hill equation fits, we estimate dissociation constant (*K*_*D*_) values for DOX binding to CB13- DNA-SWNT of 48-100 µM for peak intensity, and a *K*_*D*_ of 14-28 µM for peak wavelength. The limit of detection was calculated using 3**σ**_*SD,baseline*_ of a nanosensor control measurement with the fits for intensity and peak wavelength to obtain DOX limit of detection values of 9 µM and 8 µM, respectively. The sensitivity of the CB13-DNA-SWNT nanosensor therefore approaches peak circulatory concentrations of DOX for common therapeutic administration^30^ and could reveal areas of overly toxic drug concentrations or drug accumulation in tissue by direct imaging. Importantly, we observe a negligible fluorescence modulation of CB13-DNA-SWNT upon exposure to the alkylating-agent dacarbazine (DTIC), a chemotherapeutic commonly co-injected with DOX, which demonstrates the specificity of the nanosensor over this interfering analyte (**Figure S7**).

We lastly evaluate the CB13-DNA-SWNT nanosensor for DOX imaging in tissue, to assess feasibility of nanosensor applications in direct monitoring of DOX accumulation in biological tissues which could circumvent the use of invasive tissue biopsies.^13,31,32^ The CB13-DNA-SWNT nanosensor, prepared using the optimized SWNT:DNA ratio 3:0.8, was incorporated into muscle tissue excised from the hind leg of a mouse by incubating a 10 mg/L concentration of CB13-DNA-SWNT with the tissue. Near-infrared images of the muscle tissue after incubation and washing of CB13-DNA-SWNTs not associated with the tissue reveal clear localization of nanosensors within the striated tissue (**Figure 6**). Addition of 100 µM DOX to the saline bath containing the nanosensor-loaded tissue caused a pronounced red-shift of the nanosensor fluorescence emission spectrum integrated across the entire field of view. The 7 nm nanosensor red-shift signal was observed within 20 min of DOX addition to the saline bath. The nanosensor red-shift was accompanied by a decrease in fluorescence intensity, in contrast to the increase observed *in vitro* using the same nanosensor sample. This discrepancy is likely a result of varying ionic conditions and adsorption of proteins from the tissue, which are known to attenuate SWNT fluorescence intensity.^33,34^ Our results demonstrate that SWNT fluorescence wavelength shifts provide a robust optical readout for chemotherapeutic drug imaging in the relevant biological milieu of muscle tissue.^5,35^

**Figure 6.**
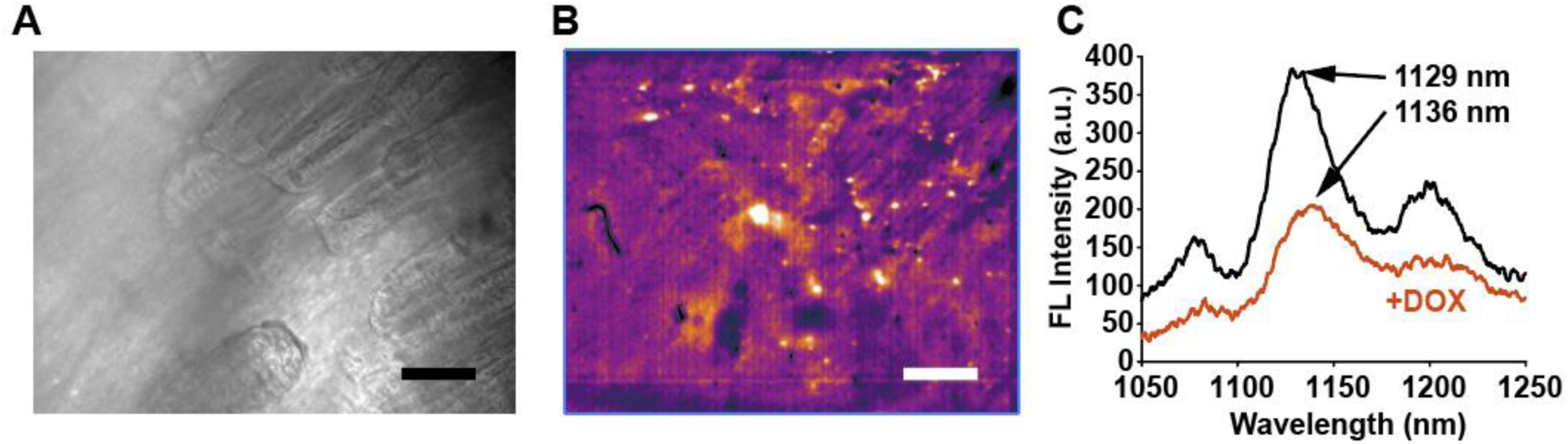
(**A**) Brightfield image of mouse muscle excised from the upper hind leg. (**B**) NIR image of muscle tissue from the same field of view as **A**, revealing fluorescence from CB13-DNA-SWNT localized in the muscle tissue. Scale bar is 100 µm. (**C**) Emission spectra collected from (**B**) before (black) and after (red) addition of 100 µM DOX to the sample bath.

## Conclusions

This work presents a chemometric approach for analyzing NIR emission spectra obtained from library screens of SWNT nanosensors, and identifying SWNT nanosensors with desired fluorescence signals. These methods provide both a rapid, quantitative measure for evaluating potential new nanosensors for biochemical analytes, and a chemometric platform to infer relationships between analyte structure and nanosensor response. Such approaches are increasingly important for nanosensor discovery and understanding interaction mechanisms as high-throughput compatible screening methods become increasingly available. Moreover, we show that application of these methods can successfully identify a new candidate optical nanosensor for the chemotherapeutic DOX, with a wavelength-shifting optical readout that is compatible with imaging in the biological milieu of muscle tissue. We show that changes to both the intensity and wavelength of our near-infrared nanosensor in response to DOX, and its selectivity over co-administered chemotherapeutic dacarbazine, enables imaging of DOX in mouse hindlimb tissue.

Methods for the continuous, real-time measurements of drugs *in vivo* are necessary to facilitate the development of personalized therapies that employ feedback-controlled delivery to improve safety and efficacy. While great strides have been made in monitoring circulatory concentrations of small molecule drugs with high sensitivities and speeds using electrochemical sensors,^36^ recording spatial distribution and accumulation in tissue remains a challenge. Visible wavelength fluorophores lack the tissue-penetrating ability required for *in vivo* monitoring, making tissue biopsy a necessary step for quantifying drugs and metabolites in patients. As such, the approach we demonstrate herein may be leveraged for future generations of SWNT-based nanosensors towards non-invasive, deep-tissue chronic imaging of drug distribution and accumulation.

## Acknowledgements

The authors thank A Beyene and D Yang for providing samples. We acknowledge funding from Burroughs Wellcome Fund Career Award at the Scientific Interface (CASI) (M.P.L), a Stanley Fahn PDF Junior Faculty Grant with Award # PF-JFA-1760 (M.P.L), a Beckman Foundation Young Investigator Award (M.P.L), a FFAR New Innovator Award, a NIH NIDA CEBRA award # R21DA044010 (to M.P.L.), and a DARPA Young Investigator Award (M.P.L). M.P.L. is a Chan Zuckerberg Biohub investigator. J.T.D.O. is supported by an Early Investigator Research Award from the Congressionally Directed Medical Research Program through the US Department of Defense. R.P. is supported by an NSF Graduate Research Fellowship. J.T.D.O. and M.P.L. both received support from a SAS Institute-Burroughs Wellcome Fund Collaborative Fellowship.

## References

(1) You, L., Zha, D., and Anslyn, E. V. (2015) Recent Advances in Supramolecular Analytical Chemistry Using Optical Sensing. Chem. Rev. 115, 7840–7892.

(2) Jin, H., Heller, D. A., Kalbacova, M., Kim, J.-H., Zhang, J., Boghossian, A. A., Maheshri, N., and Strano, M. S. (2010) Detection of single-molecule H2O2 signalling from epidermal growth factor receptor using fluorescent single-walled carbon nanotubes. Nat. Nanotechnol. 5, 302–309.

(3) Zhang, J., Landry, M. P., Barone, P. W., Kim, J. H., Lin, S., Ulissi, Z. W., Lin, D., Mu, B., Boghossian, A. A., Hilmer, A. J., Rwei, A., Hinckley, A. C., Kruss, S., Shandell, M. A., Nair, N., Blake, S., Şen, F., Şen, S., Croy, R. G., Li, D., Yum, K., Ahn, J. H., Jin, H., Heller, D. A., Essigmann, J. M., Blankschtein, D., and Strano, M. S. (2013) Molecular recognition using corona phase complexes made of synthetic polymers adsorbed on carbon nanotubes. Nat. Nanotechnol. 8, 959–968.

(4) Landry, M. P., Ando, H., Chen, A. Y., Cao, J., Kottadiel, V. I., Chio, L., Yang, D., Dong, J., Lu, T. K., and Strano, M. S. (2017) Single-molecule detection of protein efflux from microorganisms using fluorescent single-walled carbon nanotube sensor arrays. Nat. Nanotechnol. 12, 368–377.

(5) Harvey, J. D., Jena, P. V., Baker, H. A., Zerze, G. H., Williams, R. M., Galassi, T. V., Roxbury, D., Mittal, J., and Heller, D. A. (2017) A carbon nanotube reporter of microRNA hybridization events in vivo. Nat. Biomed. Eng. 1, 0041.

(6) Bisker, G., Ahn, J., Kruss, S., Ulissi, Z. W., Salem, D. P., and Strano, M. S. (2015) A Mathematical Formulation and Solution of the CoPhMoRe Inverse Problem for Helically Wrapping Polymer Corona Phases on Cylindrical Substrates. J. Phys. Chem. C 119, 13876–13886.

(7) Ulissi, Z. W., Sen, F., Gong, X., Sen, S., Iverson, N., Boghossian, A. A., Godoy, L. C., Wogan, G. N., Mukhopadhyay, D., and Strano, M. S. (2014) Spatiotemporal Intracellular Nitric Oxide Signaling Captured Using Internalized, Near-Infrared Fluorescent Carbon Nanotube Nanosensors. Nano Lett. 14, 4887–4894.

(8) Welsher, K., Sherlock, S. P., and Dai, H. (2011) Deep-tissue anatomical imaging of mice using carbon nanotube fluorophores in the second near-infrared window. Proc. Natl. Acad. Sci. U. S. A. 108, 8943–8.

(9) Del Bonis-O’Donnell, J. T., Page, R. H., Beyene, A. G., Tindall, E. G., McFarlane, I. R., and Landry, M. P. (2017) Dual Near-Infrared Two-Photon Microscopy for Deep-Tissue Dopamine Nanosensor Imaging. Adv. Funct. Mater. 27, 1702112.

(10) Salem, D. P., Landry, M. P., Bisker, G., Ahn, J., Kruss, S., and Strano, M. S. (2016) Chirality dependent corona phase molecular recognition of DNA-wrapped carbon nanotubes. Carbon N. Y. 97, 147–153.

(11) Heller, D. A., Jin, H., Martinez, B. M., Patel, D., Miller, B. M., Yeung, T. K., Jena, P. V., Hobartner, C., Ha, T., Silverman, S. K., and Strano, M. S. (2009) Multimodal optical sensing and analyte specificity using single-walled carbon nanotubes. Nat. Nanotechnol. 4, 114–120.

(12) Chatterjee, K., Zhang, J., Honbo, N., and Karliner, J. S. (2010) Doxorubicin cardiomyopathy. Cardiology 115, 155–62.

(13) Rahman, A. M., Yusuf, S. W., and Ewer, M. S. (2007) Anthracycline-induced cardiotoxicity and the cardiac-sparing effect of liposomal formulation. Int. J. Nanomedicine 2, 567–83.

(14) Kruss, S., Landry, M. P., Vander Ende, E., Lima, B. M. A., Reuel, N. F., Zhang, J., Nelson, J., Mu, B., Hilmer, A., and Strano, M. (2014) Neurotransmitter detection using corona phase molecular recognition on fluorescent single-walled carbon nanotube sensors. J. Am. Chem. Soc. 136, 713–724.

(15) Bro, R., and Smilde, A. K. (2014) Principal component analysis. Anal. Methods 6, 2812–2831.

(16) Stewart, S., Ivy, M. A., and Anslyn, E. V. (2014) The use of principal component analysis and discriminant analysis in differential sensing routines. Chem. Soc. Rev. 43, 70–84.

(17) Jaumot, J., Eritja, R., Navea, S., and Gargallo, R. (2009) Classification of nucleic acids structures by means of the chemometric analysis of circular dichroism spectra. Anal. Chim. Acta 642, 117–126.

(18) Polo, E., and Kruss, S. (2016) Impact of Redox-Active Molecules on the Fluorescence of Polymer-Wrapped Carbon Nanotubes. J. Phys. Chem. C 120, 3061–3070.

(19) Mark, H. B., Atta, N., Ma, Y., Petticrew, K., Zimmer, H., Lunsford, S., Rubinson, J., and Galal, A. (1995) The electrochemistry of neurotransmitters at conducting organic polymer electrodes: electrocatalysis and analytical applications. Bioelectrochemistry Bioenerg. 38, 229–245.

(20) Das, A., Sood, A. K., Maiti, P. K., Das, M., Varadarajan, R., and Rao, C. N. R. (2008) Binding of nucleobases with single-walled carbon nanotubes: Theory and experiment. Chem. Phys. Lett. 453, 266–273.

(21) Sarmah, A., and Roy, R. K. (2013) Understanding the Interaction of Nucleobases with Chiral Semiconducting Single-Walled Carbon Nanotubes: An Alternative Theoretical Approach Based on Density Functional Reactivity Theory. J. Phys. Chem. C 117, 21539–21550.

(22) Strano, M. S., Huffman, C. B., Moore, V. C., O’Connell, M. J., Haroz, E. H., Hubbard, J., Miller, M., Rialon, K., Kittrell, C., Ramesh, S., Hauge, R. H., and Smalley, R. E. (2003) Reversible, Band-Gap-Selective Protonation of Single-Walled Carbon Nanotubes in Solution. J. Phys. Chem. B 107, 6979–6985.

(23) Dukovic, G., White, B. E., Zhou, Z., Wang, F., Jockusch, S., Steigerwald, M. L., Heinz, T. F., Friesner, R. A., Turro, N. J., and Brus, L. E. (2004) Reversible surface oxidation and efficient luminescence quenching in semiconductor single-wall carbon nanotubes. J. Am. Chem. Soc. 126, 15269–15276.

(24) Salem, D. P., Gong, X., Liu, A. T., Koman, V. B., Dong, J., and Strano, M. S. (2017) Ionic Strength-Mediated Phase Transitions of Surface-Adsorbed DNA on Single-Walled Carbon Nanotubes. J. Am. Chem. Soc. 139, 16791–16802.

(25) Liu, Z., Fan, A. C., Rakhra, K., Sherlock, S., Goodwin, A., Chen, X., Yang, Q., Felsher, D. W., and Dai, H. (2009) Supramolecular stacking of doxorubicin on carbon nanotubes for in vivo cancer therapy. Angew. Chemie - Int. Ed. 48, 7668–7672.

(26) Choi, J. H., and Strano, M. S. (2007) Solvatochromism in single-walled carbon nanotubes. Appl. Phys. Lett. 90, 223114.

(27) Heller, D. A., Jeng, E. S., Yeung, T.-K., Martinez, B. M., Moll, A. E., Gastala, J. B., and Strano, M. S. (2006) Optical detection of DNA conformational polymorphism on single-walled carbon nanotubes. Science 311, 508–11.

(28) Harvey, J. D., Baker, H. A., Mercer, E., Budhathoki-Uprety, J., and Heller, D. A. (2017) Control of Carbon Nanotube Solvatochromic Response to Chemotherapeutic Agents. ACS Appl. Mater. Interfaces 9, 37947–37953.

(29) Beyene, A. G., Alizadehmojarad, A. A., Dorlhiac, G., Streets, A. M., Kral, P., Vuković, L., and Landry, M. P. (2018) Ultralarge Modulation of Single Wall Carbon Nanotube Fluorescence Mediated by Neuromodulators Adsorbed on Arrays of Oligonucleotide Rings. bioRxiv 351627.

(30) Barpe, D. R., Rosa, D. D., and Froehlich, P. E. (2010) Pharmacokinetic evaluation of doxorubicin plasma levels in normal and overweight patients with breast cancer and simulation of dose adjustment by different indexes of body mass. Eur. J. Pharm. Sci. 41, 458–463.

(31) Iverson, N. M., Barone, P. W., Shandell, M., Trudel, L. J., Sen, S., Sen, F., Ivanov, V., Atolia, E., Farias, E., McNicholas, T. P., Reuel, N., Parry, N. M. A., Wogan, G. N., and Strano, M. S. (2013) In vivo biosensing via tissue-localizable near-infrared-fluorescent single-walled carbon nanotubes. Nat. Nanotechnol. 8, 873–880.

(32) Stewart, D. J., Grewaal, D., Green, R. M., Mikhael, N., Goel, R., Montpetit, V. A., and Redmond, M. D. (1993) Concentrations of doxorubicin and its metabolites in human autopsy heart and other tissues. Anticancer Res. 13, 1945–52.

(33) Gong, X., Sharma, A. K., Strano, M. S., and Mukhopadhyay, D. (2014) Selective Assembly of DNA-Conjugated Single-Walled Carbon Nanotubes from the Vascular Secretome. ACS Nano 8, 9126–9136.

(34) Ge, C., Du, J., Zhao, L., Wang, L., Liu, Y., Li, D., Yang, Y., Zhou, R., Zhao, Y., Chai, Z., and Chen, C. (2011) Binding of blood proteins to carbon nanotubes reduces cytotoxicity. Proc. Natl. Acad. Sci. U. S. A. 108, 16968–73.

(35) Jena, P. V., Roxbury, D., Galassi, T. V., Akkari, L., Horoszko, C. P., Iaea, D. B., Budhathoki-Uprety, J., Pipalia, N., Haka, A. S., Harvey, J. D., Mittal, J., Maxfield, F. R., Joyce, J. A., and Heller, D. A. (2017) A Carbon Nanotube Optical Reporter Maps Endolysosomal Lipid Flux. ACS Nano 11, 10689–10703.

(36) Arroyo-Currás, N., Somerson, J., Vieira, P. A., Ploense, K. L., Kippin, T. E., and Plaxco, K. W. (2017) Real-time measurement of small molecules directly in awake, ambulatory animals. Proc. Natl. Acad. Sci. 114, 645–650.

